# Preprints vs. Journal Articles: Citation Impact in COVID-19 Research

**DOI:** 10.1101/2024.11.08.622753

**Authors:** Hiroyuki Tsunoda, Yuan Sun, Masaki Nishizawa, Xiaomin Liu, Kou Amano, Rie Kominami

## Abstract

Focusing on article and citation count, this study examined 110 academic journals whose papers on COVID-19 were indexed in PubMed from 2020 to 2023. To analyze these journals’ characteristics, this study introduced the concepts of paper ratio, citation ratio, and citation impact. Journals with similar trends in paper and citation count were positioned near the diagonal on a two-dimensional graph (y=x) with paper ratio on the x-axis and citation ratio on the y-axis, indicating a slope close to 1. Conversely, the greater the disparity between paper ratio and citation ratio, the wider the angle between the journal’s position and the diagonal. Of the 110 journals in 2020, 83 were above the diagonal with a citation impact greater than 1, while 27 were below the diagonal with a citation impact of 1 or lower. Notably, nine journals—*BMC Health Services Research, Genome Medicine, Nature, Annals of Epidemiology, BMC Bioinformatics, Nature Microbiology, PLoS Computational Biology, Medicine (Baltimore)*, and *Journal of Clinical Investigation*—showed high citation impact, with preprint-distributed papers being cited nearly five times more than directly submitted papers. This study demonstrates that articles initially distributed as preprints in these journals tend to receive substantially more citations than directly submitted ones.

## 1 INTRODUCTION

A preprint is a scientific paper that has yet to undergo peer review for publication in an academic journal but can be nevertheless viewed and downloaded from an archive and may also be cited. The COVID-19 pandemic saw an increase in the value and importance of preprints as they helped to disseminate public health information. On January 30, 2020, the World Health Organization (WHO) declared COVID-19 a public health emergency of international concern and, on March 11, 2020, declared it a pandemic because of the global spread and severity of the infection. Since 2020, preprints have served as a new means of academic communication [1, 7, 22]. Although preprints have substantially increased in number across various fields, their use continues to be limited to certain regions and subjects [21]. One reason for authors’ hesitancy is their concern regarding the public posting of non-peer-reviewed research. Many researchers are also unaware that preprints [6]. While some have argued that non-peer-reviewed preprints have quality issues, research has shown that their contents are generally consistent with those of the final published versions after peer review [2, 3, 9, 10]. In addition, many preprints are eventually published in respected journals, a testament that their quality is not inferior [13]. Furthermore, papers initially distributed as preprints often undergo faster peer review upon their submission to journals, leading to quicker publication [14, 15]. Compared with non-COVID-19 papers, COVID-19 articles are accepted more frequently and complete peer review more quickly [5, 11, 16, 17]. Preprints have also received considerable media coverage [4, 20] and have been widely disseminated on social media [12], which may explain their higher post-peer-review citation rates. In medicine, preprints on COVID-19 tend to be accepted by high-impact journals [18]. Investigations into papers with COVID-19 Medical Subject Headings (MeSH) in PubMed have revealed that these studies receive more citations than any other journal articles. The early preprint distribution of papers on COVID-19 has resulted in high citation counts [8, 19]. This study seeks to identify and clarify the characteristics of academic journals containing articles on COVID-19 that are distributed as preprints and have received substantially more citations than those that have been directly submitted.

## 2 DATA AND METHOD

### 2.1 Screening of Preprint-Distributed Papers

This study used bioRxiv’s application programming interface (API) to extract and download metadata for journal articles published after being distributed as preprints. The API endpoint format is https://api.biorxiv.org/pubs/[server]/[interval]/[cursor]. We set the server parameter to either bioRxiv or medRxiv as the preprint source.

### 2.2 Journal Articles on COVID-19

PubMed is a free database that allows for the search and retrieval of biomedicine and life sciences studies to improve global and individual health. Available to the public online since 1996 and containing about 37 million citations and abstracts of biomedical literature, PubMed was developed and is maintained by the National Center for Biotechnology Information (NCBI) at the U.S. National Library of Medicine (NLM), which is under the National Institutes of Health (NIH). For a comprehensive collection of COVID-19 articles, this study gathered papers published on PubMed. To this end, we used Entrez Programming Utilities (E-utilities), a publicly accessible API for the NCBI Entrez system written in Python. The subjects of the papers were determined using the NLM’s MeSH thesaurus. Papers tagged with “COVID-19” in MeSH were classified as COVID-19 papers, while those not tagged with this term were considered non-COVID-19 articles.

### 2.3 Collection Period, Types of Documents, and Citation Counts

Because academic journals have already contained papers on COVID-19 topics in 2019, the collection period for this study was set to 2019–2023. However, because of the small number of papers published in 2019, the actual analysis focused on the period 2020–2023. There were various types of documents published in academic journals; this study focused on articles and reviews to measure impact using citation counts. Although researchers commonly use Clarivate’s Times Cited from Web of Science to measure impact in citation analysis, its restrictions on programmatic downloading necessitated the use of PubMed in this study, which allows for citation downloads via API, where a paper’s citation count is determined by counting the articles listed in the “Cited by” section, and the metadata for all analyzed studies was downloaded from PubMed in June 2024.

### 2.4 Paper Ratio, Citation Ratio, and Citation Impact

We propose the following three indicators to examine papers on a specific topic published in journals, the citation patterns they received, and the influence of the topic on these citations. The paper ratios, citation ratios, and citation impact values of sets A (journal articles) and set B (journal articles that cite set A) are defined as follows. Paper ratio refers to the number of elements in set A divided by the number of elements in its complement (equation 1).

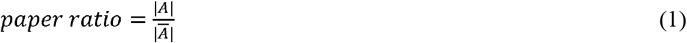

Citation ratio pertains to the number of elements in set B divided by the number of elements in its complement (equation 2).

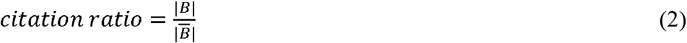

Citation impact refers to the ratio of the citation ratio to the paper ratio (equation 3).

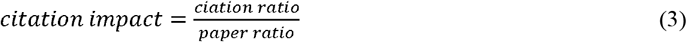

### 2.5 Journal Selection

Between 2020 and 2023, a total of 183 journals published at least 10 PubMed-indexed preprint-distributed papers on COVID-19. However, our analysis focused on only 110 journals that contained complete data for all years.

## ANALYSIS AND RESULTS

### 3.1 Tendencies of COVID-19 Papers to Receive Citations

Table 1 presents the citation-receiving statuses of preprint-distributed COVID-19 and non-COVID-19 papers. It shows that the 110 analyzed journals in 2020 had a median paper ratio of 0.25 for COVID-19 preprint-distributed studies and a median citation ratio of 0.38. In contrast, a median paper ratio of 0.03 was observed for non-COVID-19 preprint-distributed papers along with a median citation ratio of 0.05. COVID-19 papers had a citation impact of 1.52, while the value for non-COVID-19 papers was 1.42, with a difference of 0.10 (1.52 - 1.42), indicating that the former set received citations more efficiently. The difference in citation impact continued to show the same trend through 2023.

**Table 1:**
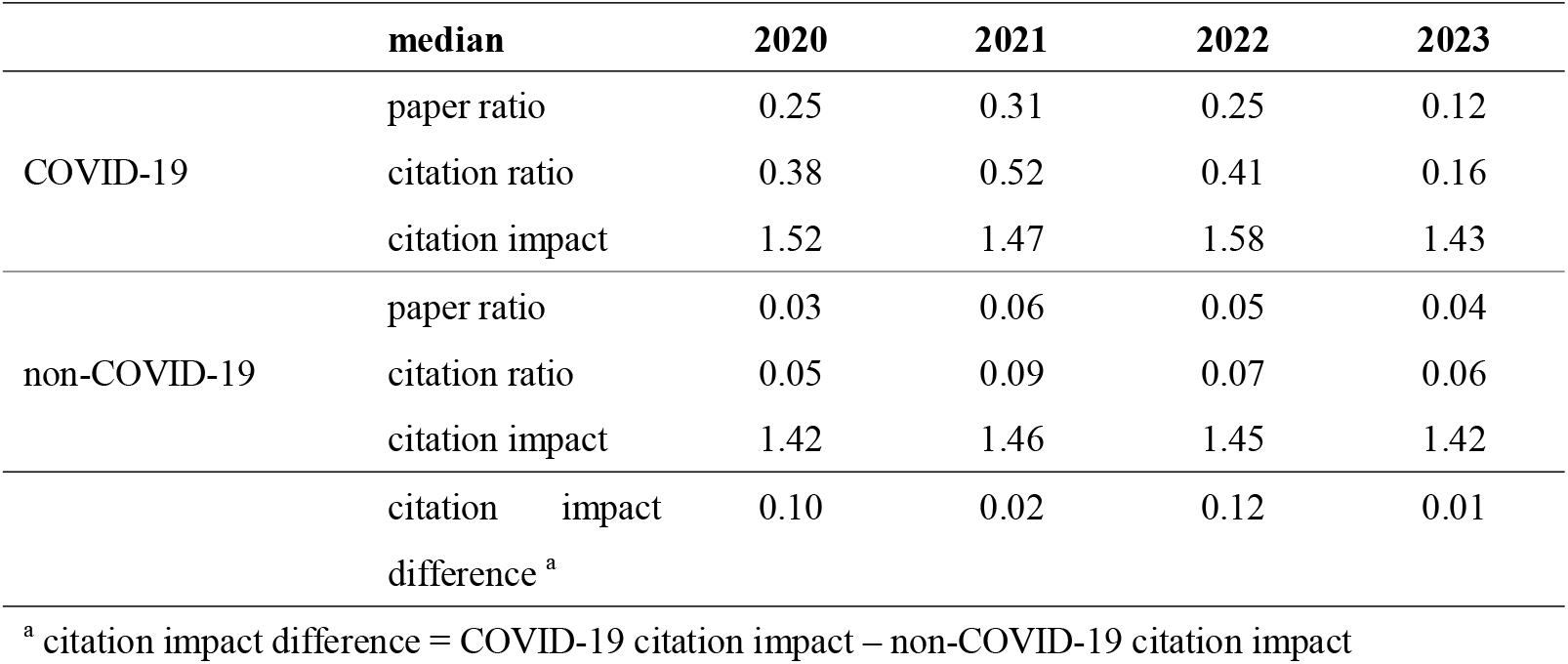
Median Paper Ratio, Median Citation Ratio, and Median Citation Impact for Preprint-Distributed and Directly Submitted COVID-19 and non-COVID-19 papers (2020–2023)

|  | median | 2020 | 2021 | 2022 | 2023 |
| --- | --- | --- | --- | --- | --- |
| COVID-19 | paper ratio | 0.25 | 0.31 | 0.25 | 0.12 |
|  | citation ratio | 0.38 | 0.52 | 0.41 | 0.16 |
|  | citation impact | 1.52 | 1.47 | 1.58 | 1.43 |
| non-COVID-19 | paper ratio | 0.03 | 0.06 | 0.05 | 0.04 |
|  | citation ratio | 0.05 | 0.09 | 0.07 | 0.06 |
|  | citation impact | 1.42 | 1.46 | 1.45 | 1.42 |
|  | citation impact difference <sup>a</sup> | 0.10 | 0.02 | 0.12 | 0.01 |

<sup>a</sup> citation impact difference = COVID-19 citation impact – non-COVID-19 citation impact

### 3.2 Paper Ratio and Citation Ratio Distribution

AS shown in Figure 1, the interquartile range (IQR) of journal distribution based on citation ratio for COVID-19 papers across all years extends further upward compared with the distribution based on paper ratio, and this discrepancy is particularly notable in 2020. One plausible explanation for this is the smaller number of published outputs in 2020, when COVID-19 research was still in its nascent stages, and papers that were initially disseminated as preprints were subsequently published, attracting considerable attention from the research community. This early-stage distribution as preprints may have contributed to the heightened citation ratios, reflecting the urgency and widespread interest in COVID-19 findings at that time. To further examine this phenomenon, we performed a comparative analysis of preprint-distributed papers and directly submitted papers on COVID-19 in 2020.

**Figure 1:**
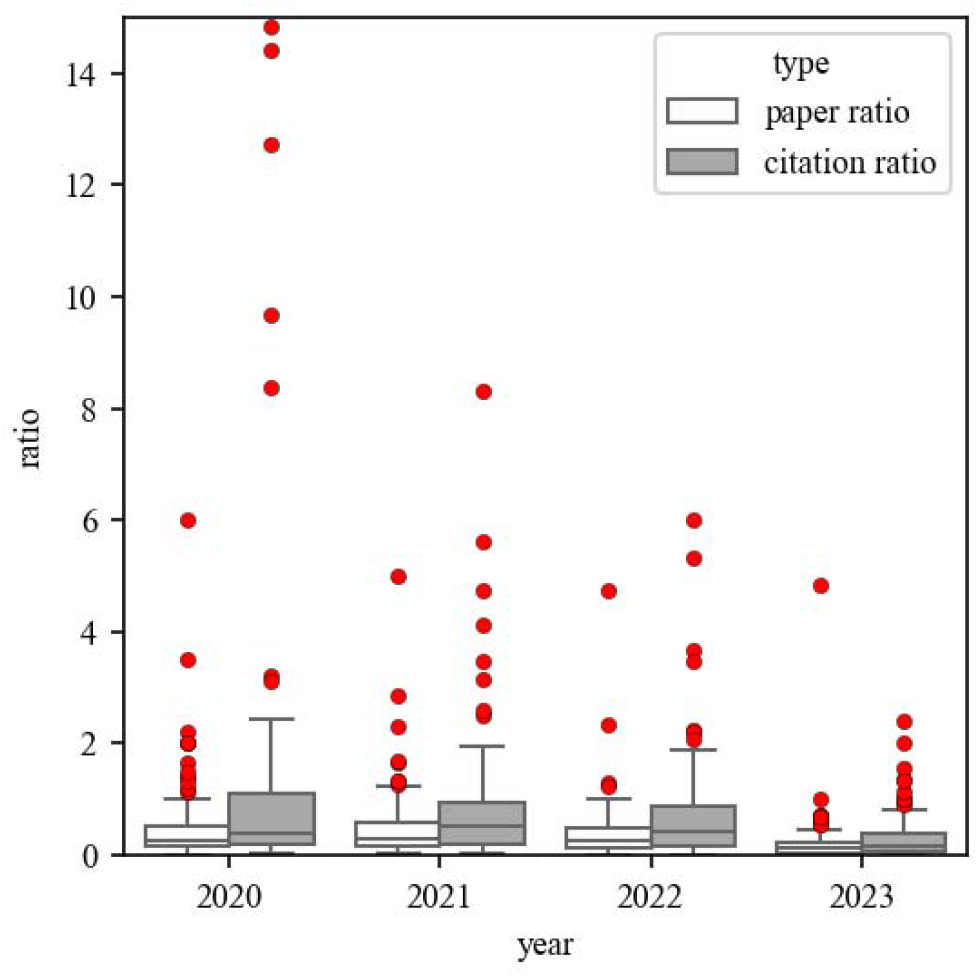
Range of Distribution for Paper Ratio and Citation Ratio of Preprint-Distributed and Directly Submitted Journal Articles Related on COVID-19 (2020–2023)

### 3.3 Distribution and Identification of High-Citation-Impact Journals in 2020

Overall, academic journals showing similar trends in paper and citation count tend to be located near the diagonal in a two-dimensional graph (y = x), where paper ratio is plotted on the x-axis and the citation ratio on the y-axis, indicating a slope close to 1. Conversely, the higher the divergence of paper and citation ratio trends, the higher the increase in the angle between the journal’s position and the diagonal line. Figure 2 illustrates the citation trends for the 110 journals in 2020, with paper ratio on the x-axis and citation ratio on the y-axis. Among these, 83 journals containing preprint-distributed articles that tend to receive more citations are represented as red and orange circles above the diagonal line (y = x). A further 9 journals showing extremely high citation impact and indicated as upper outliers in the box plot (the value is calculated by adding 1.5 times the IQR to the third quartile (Q3); for example, in 2020, the IQR was 1.4980, and the Q3 was 2.5103, resulting in an approximate upper outlier value of slightly below 4.76), have their titles displayed on the scatterplot. Conversely, 27 journals containing directly submitted articles with higher citation ratios are depicted as cyan circles below the diagonal line, but no journals were identified as lower outliers. The dotted (y = 4.76x) and diagonal line (y = x) serve as boundaries for these journals, with the dotted line specifically marking the threshold for exceptionally high citation impact. Because many circles are concentrated near the origin (0,0), a magnified view is shown in the bottom-right corner.

**Figure 2:**
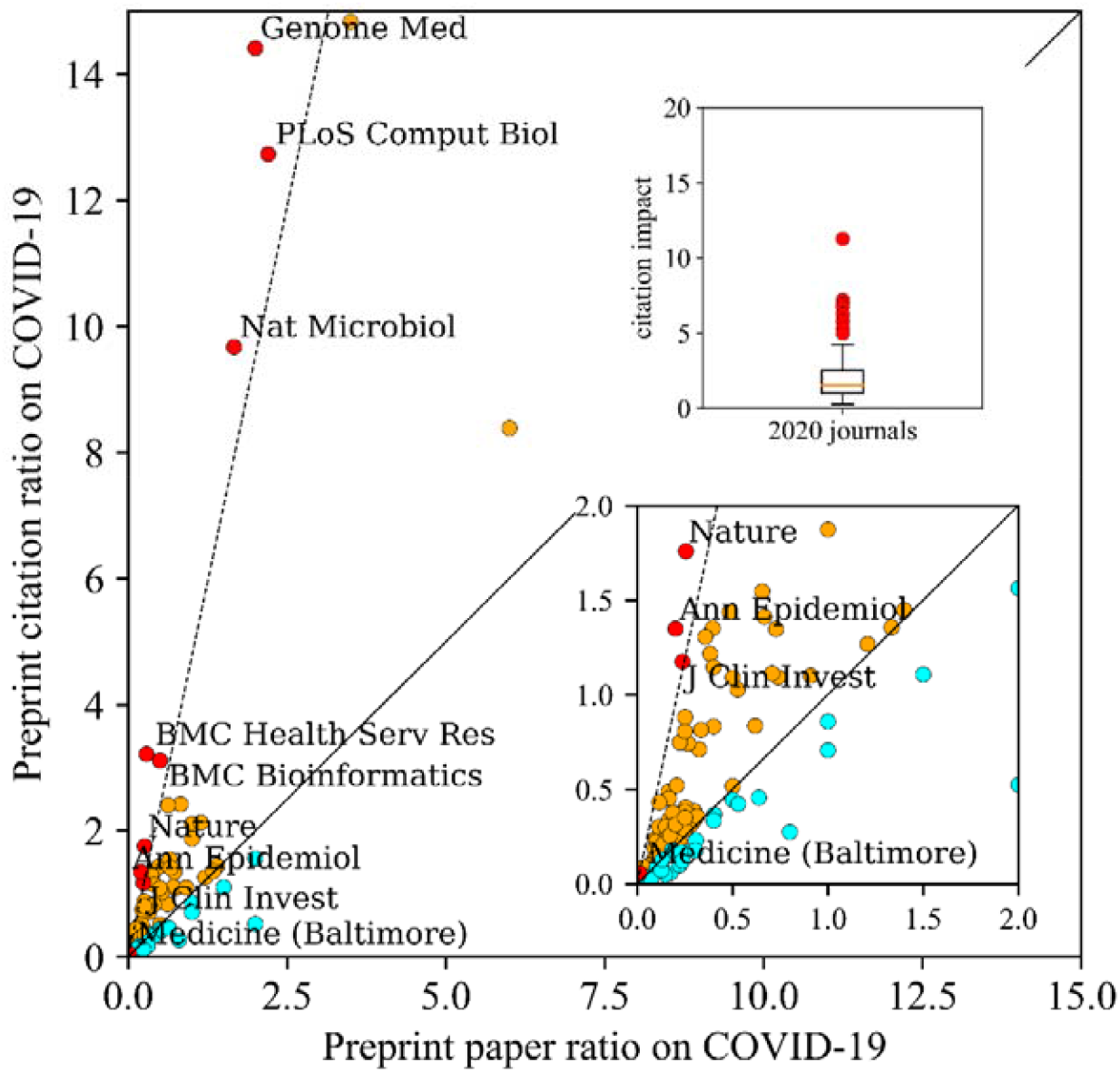
Distribution and Identification of High-Citation-Impact Journals in 2020 (Including Box Plot and Enlarged View for Low-Ratio Journals)

### 3.4 Characteristics of High-Citation-Impact Journals in 2020

Table 2 presents the top 9 out of 110 journals analyzed for 2020 ranked by strength of citation impact. The leading journal, *BMC Health Services Research*, contained two preprint-distributed papers receiving a total of 251 citations, resulting in an average of 125.5 citations per paper. In contrast, the 7 directly submitted papers in the same journal received 78 citations, with an average of 11.14 per paper. Therefore, the journal’s overall citation impact was approximately 11.26 [(251/2) / (78/7)], indicating that preprint-distributed papers attracted considerable research attention. This finding underlines how preprints can play a key role in amplifying the visibility and citation performance of early COVID-19 research. Following this, *Genome Medicine, Nature, Annals of Epidemiology, BMC Bioinformatics, Nature Microbiology, PLoS Computational Biology, Medicine (Baltimore)*, and *Journal of Clinical Investigation* all showed citation impacts of 4.76 or higher, suggesting that when the COVID-19 papers in these journals were distributed as preprints before being formally published, they received more than 4.76 times the citations compared with directly submitted papers.

**Table 2:**
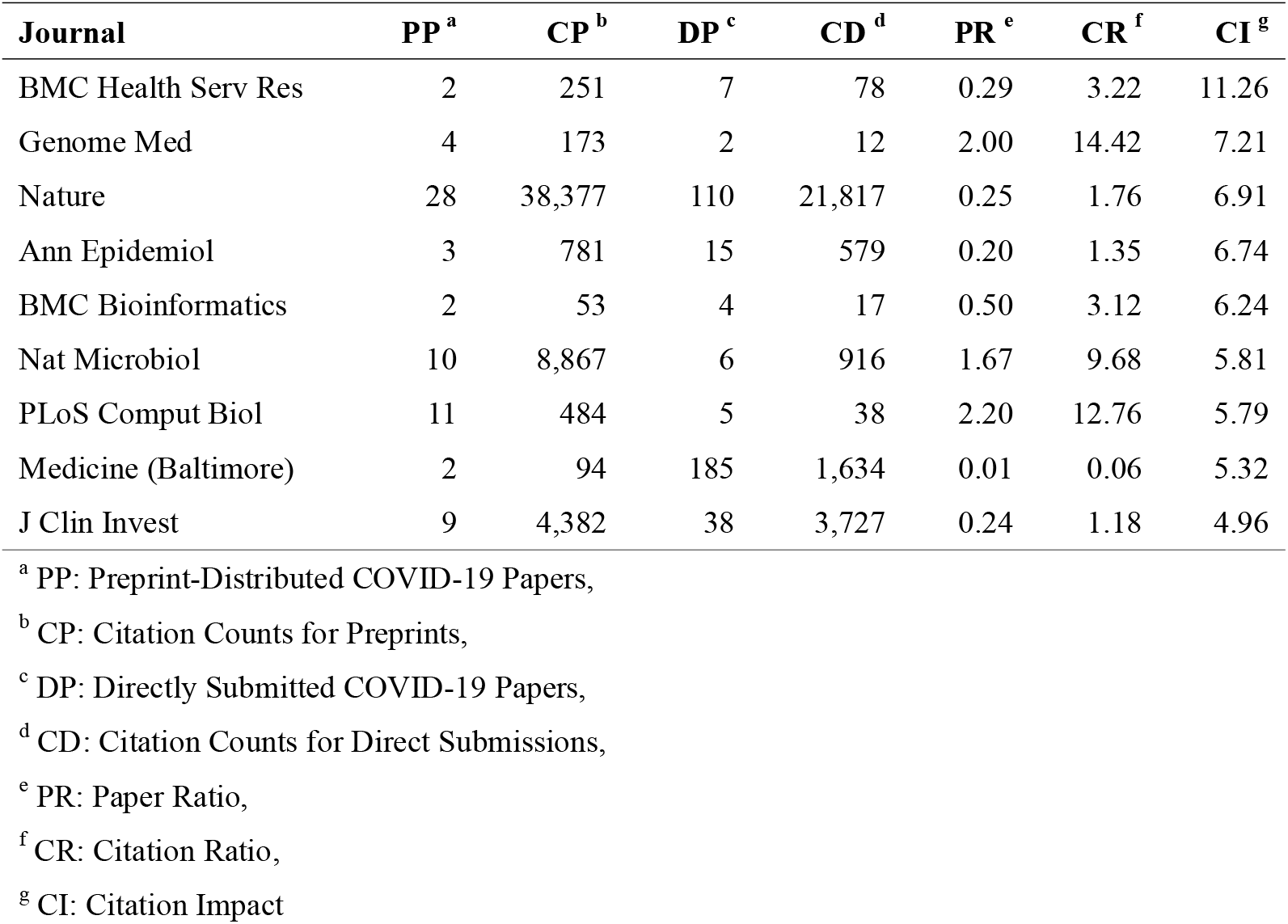
Top 9 Journals by Citation Impact in 2020.

| Journal | PP<sup>a</sup> | CP<sup>b</sup> | DP<sup>c</sup> | CD<sup>d</sup> | PR<sup>e</sup> | CR<sup>f</sup> | CI<sup>g</sup> |
| --- | --- | --- | --- | --- | --- | --- | --- |
| BMC Health Serv Res | 2 | 251 | 7 | 78 | 0.29 | 3.22 | 11.26 |
| Genome Med | 4 | 173 | 2 | 12 | 2.00 | 14.42 | 7.21 |
| Nature | 28 | 38,377 | 110 | 21,817 | 0.25 | 1.76 | 6.91 |
| Ann Epidemiol | 3 | 781 | 15 | 579 | 0.20 | 1.35 | 6.74 |
| BMC Bioinformatics | 2 | 53 | 4 | 17 | 0.50 | 3.12 | 6.24 |
| Nat Microbiol | 10 | 8,867 | 6 | 916 | 1.67 | 9.68 | 5.81 |
| PLoS Comput Biol | 11 | 484 | 5 | 38 | 2.20 | 12.76 | 5.79 |
| Medicine (Baltimore) | 2 | 94 | 185 | 1,634 | 0.01 | 0.06 | 5.32 |
| J Clin Invest | 9 | 4,382 | 38 | 3,727 | 0.24 | 1.18 | 4.96 |

<sup>a</sup> PP: Preprint-Distributed COVID-19 Papers,
<sup>b</sup> CP: Citation Counts for Preprints,
<sup>c</sup> DP: Directly Submitted COVID-19 Papers,
<sup>d</sup> CD: Citation Counts for Direct Submissions,
<sup>e</sup> PR: Paper Ratio,
<sup>f</sup> CR: Citation Ratio,
<sup>g</sup> CI: Citation Impact

## 4 CONCLUSIONS AND DISCUSSIONS

This study investigated COVID-19 papers in 110 academic journals registered in PubMed between 2020 and 2023, focusing on papers and citation counts. The characteristics of these journals were examined through the concepts of paper ratio, citation ratio, and citation impact. In 2020, a total of 83 journals whose preprint-distributed papers had a stronger tendency to acquire citations were positioned above the diagonal line (y = x) whereas 27 journals in which directly submitted papers were more prominent were located below the line. Furthermore, 9 of the 83 journals demonstrated exceptionally high citation impact values surpassing 4.76. These findings underscore the significant influence of preprints in accelerating the dissemination and visibility of COVID-19 research during a critical period. The swift citation accumulation suggests that preprints served as an essential tool for timely knowledge sharing when the demand for reliable information was particularly high.

Result showed that COVID-19 papers quickly received a considerable number of citations, attracting substantial attention from the academic community. COVID-19 papers had a substantially higher median citation count than non-COVID-19 papers for the same period. As the pandemic subsided, these papers’ citation frequency declined year by year, reflecting a shift from intense to diminished scientific research activity. This trend suggests that while preprints were particularly valuable during the peak of the pandemic, their role evolved as the urgency for new information lessened.

High-impact-factor journals published papers that were cited more frequently, indicating that scholars prioritize influential journals in their fields. This study further observed that COVID-19 studies cited early preprint-distributed papers more frequently than direct-submission papers, highlighting the important role play by preprints in academic communication during the COVID-19 pandemic and establishing a new model for scholarly exchange. The preference for preprints during the pandemic points to a broader shift in how academic communication adapts in response to global crises. Understanding the mechanisms that drove this shift could inform future strategies for rapid research dissemination. However, a more comprehensive understanding of preprints’ relatively high citation rates requires further detailed analysis, which will clarify the role of preprints in academic communication. To this end, we aim to collect and examine more data to determine how preprints influence the academic community.

## ACKNOWLEDGMENTS

This work was supported by JSPS KAKENHI grant numbers JP19K12707, JP20K12569, and JP22K12737 and ROIS NII Open Collaborative Research 2023(23FS01).

